# Stepwise approach to oocyte depletion in *Sry* mutated XY female mice

**DOI:** 10.1101/146571

**Authors:** Akihiko Sakashita, Takuya Wakai, Yukiko Kawabata, Chiaki Nishimura, Yusuke Sotomaru, Tomohiro Kono

## Abstract

Little is known regarding the mechanisms underlying infertility in male-to-female sex-reversed females. To gain a better understanding of germ cell dysfunction in this condition, we produced XY^*sry*−^-females via the CRISPR/Cas9 system in C57BL/6 inbred strain mice. Mutant mice showed severe attrition of germ cells during foetal development, resulting in depletion of ovarian germ cells before sexual maturation. Comprehensive transcriptome analysis of embryonic day 13.5 primordial germ cells (PGCs) and postnatal day 1 oocytes demonstrated that XY^*sry*−^-PGCs had already deviated from the developmental process at the mitotic stage. In addition, XY^*sry*−^-oocytes resulted in meiotic disruption and developmental failure, and the genes related to these processes were significantly changed. This grievous disruption, which caused germ cell deterioration in XY^*sry*−^-females, was shown to proceed from the germ cells themselves. These results provide novel insight into the germ cell depletion of sex-reversed mice as well as into disorders of sex differentiation in human, such as ‘Swyer syndrome’, wherein patients present as typical females albeit with an XY karyotype and infertile.

## Introduction

Sex determining region Y (*Sry*), is widely conserved in mammals and acts as a determinant in the ‘bipotential’ gonads present early in development for male sex-inducing testis formation (Kashimada and Koopman, 2010; Tanaka and Nishinakamura, 2014). In contrast, XX female embryos automatically develop into females, forming ovaries chiefly of oocytes and follicle cells (Suzuki et al., 2015). Failure of the sex differentiation process causes various types of genetic diseases termed disorders of sex differentiation in human (Jiang et al., 2016; Tam et al., 2016; Zhong and Layman, 2012). In mice, *Sry* expression occurs in GATA binding protein 4- and Wilms tumor 1 homolog-expressing gonadal somatic cells of the genital ridges at embryonic day (E) 11.0–12.0 (Kashimada and Koopman, 2010; Miyamoto et al., 2008). This expression directly supports gonadal cell precursors such as Sertoli cells that the fate of bipotential gonads is specified by E13.5, resulting in their subsequent differentiation into either ovaries or testes. In particular, *Sry* is expression triggers the activation of the male sex differentiation program and induces the expression of SRY-related high-mobility group box 9 (*Sox9*), and fibroblast growth factor 9, which leads to male gonad differentiation accompanied by formation of pre-spermatogonia and Sertoli cells (Kashimada and Koopman, 2010; Tanaka and Nishinakamura, 2014).

Notably, *Sry* mutations are well-known to result in male-to-female sex-reversed individuals. Such sex-reversed mice (XY-females) were first produced from chimeric mice via embryonic stem (ES) cells that carried a mutation at the testis-determining region of the Y chromosome (*Tdy*) by using a retroviral vector (Gubbay et al., 1990; Gubbay et al., 1992; Lovell-Badge and Robertson, 1990). More recently, a TALEN-mediated gene editing procedure has been used to target specific Y chromosome genes, resulting in the production of anatomical sex-reversed XY females were produced by deletion of the HMG domain of *Sry* (Kato et al., 2013; Wang et al., 2013). These animals were found to be infertile: however detailed analysis for infertility was not performed because the necessary number of XY-females was unavailable. XY female mice have also been produced by the deletion of autosomal *Sry*-related genes. For example, the deletion of *Sox9* in the gonadal somatic cells of male embryos resulted in their development into sex-reversed XY females (Lavery et al., 2011). In addition, the alteration of histone epigenetic signatures through deletion of the JmjC-containing H3K9 demethylase *Jmjd1a,* which positively regulates *Sry* expression, also resulted in male-to-female sex-reversed mice (Kuroki et al., 2013). Furthermore, it is known that male-to-female sex-reversed mice are produced when C57BL/6 mice are mated with certain strains of *Mus musculus,* such as B6-Y^AKR^, B6-Y^POS^, or B6-Y^TIR^ (Correa et al., 2012; Coward et al., 1994; Eicher et al., 1982; Lee and Taketo, 1994; Washburn et al., 2001). Specifically, these XY female mice exhibit an anatomically female phenotype with ovary formation; however, they are generally infertile, exhibiting extensive loss of germ cells in the ovary except in a few cases comprising hybrid strains (Kato et al., 2013; Lovell-Badge and Robertson, 1990; Mahadevaiah et al., 1993; Park and Taketo, 2003; Taketo, 2015; Vanderhyden et al., 1997; Vernet et al., 2014b; Wang et al., 2013). Overall, it is evident that XY females form well-constructed foetal gonads consisting of sites of cyst formation, pregranulosa cells, and oogonia. However, if oocytes from XY females are ovulated and fertilized they cease development at a very early stage (Vernet et al., 2014b; Villemure et al., 2007; Wang et al., 2013; Wong et al., 2000).

In comparison, unlike in human, XO mice are fertile (Cloutier et al., 2015; Probst et al., 2008), which indicates that genes located on the Y chromosome exert a pivotal effect on germ cell fate in XY females. Using a genetic model of XO mice carrying genes located on the short arm of the Y chromosome, the function of the Y-associated genes has been investigated (Vernet et al., 2014b). For example, *Zfy2* are shown to be essential for promoting Meiosis II (Vernet et al., 2011; Vernet et al., 2014a), whereas the testis determinant factor *Sry* and the spermatogonial proliferation factor eukaryotic translation initiation factor 2, subunit 3, structural gene Y-linked (*Eif2s3y*) were found to constitute minimum factors for producing live offspring from XO males, by ROSI (Yamauchi et al., 2014). In addition, deletion of *Eif2s3y* resulted in infertility because of a defect in spermatogenesis (Matsubara et al., 2015), whereas mutations of ubiquitously transcribed tetratricopeptide repeat gene showed no effect (Wang et al., 2013). Furthermore, every sex-reversed mouse tested was infertile; therefore, the specific molecular mechanisms underlying the generation of germ cell developmental defects in XY-female mice remained to be elucidated.

To gain a better understanding of the dysfunction of germ cells, developmental defects and infertility in male-to-female sex-reversed mice, we produced ample numbers of XY^*sry−*^-females using CRISPR/Cas9 system in the inbred C57BL/6 mouse strain, which is necessary to minimize experimental variation. Focusing on germ cell proliferation, attrition and functionality, comprehensive comparative transcriptomic analysis was conducted at E13.5, postnatal day (P) 1 and juvenile (4-weeks) stages. From these data, the gene sets specific for sex-reversed XY^*sry−*^-germ cells were identified and subjected to further bioinformatics analysis. The present study demonstrated that XY^*sry−*^-primordial germ cells (PGCs) of sex-reversed mice with Sry-mutation already had deviated from the developmental process at the meiotic stage. Furthermore, this disruption, which caused germ cell deterioration in XY^*sry−*^-females, appeared to proceed from the germ cells, themselves.

## Results

### Phenotype of *Sry* targeted mice

In total, we generated and used 65 sex-reversed XY^*sry−*^-female mice for germ cell analysis (Fig S1A), in which part of the *Sry* sequence adjacent to the transcription start site was deleted using the CRISPR/Cas9 system. The generated mice were anatomically normal females, although slight aplasia was detected in the reproductive organs (Fig. 1A). Sequence analysis detected either 1 bp-14 bp deletions or large insersions (500 bp<) in the *Sry* gene (Fig. S1B), which led to framesifts of the gene and resulted in the loss of function in all cases. Of the mutations detected in the XY^*sry−*^-female mice, 7 bp deletions (n=23.4%) most frequently occurred (Fig. S1). All mutations detected in XY^*sry−*^-females led to translational frameshifts in SRY and its functional failure. Mating tests of 14 XY^*sry−*^-females (8-10 weeks old) with normal XY males revealed the infertility of the sex-reversed mice (Fig. 1B). To determine the cause of infertility, the numbers of PGCs and oocytes were examined. Notably, a significant decrease in PGC number was already found at E13.5 in the XY^*sry−*^-females (Fig. 1C), and the oocyte numbers at P1 and P8 were markedly decreased to 1/5 and 1/6 compared with those of controls, respectively. Immunofluorescence staining of DDX4, a germ cell maker, indicated that fewer positive cells were apparent in XY females. In particular, the primary follicle oocytes were lost by the P8 stage in XY^*sry−*^-females (Fig. 1D). The decrease of PGCs in the XY^*sry−*^-female ovary progressed quickly after mid gestation, finally resulting in ovarian atrophy accompanied with oocyte depletion by 6 weeks of age (Fig. 1E). These results revealed that the infertility of sex-reversed XY^*sry−*^-female mice was caused by complete depletion of the germ cells.

**Fig. 1.**
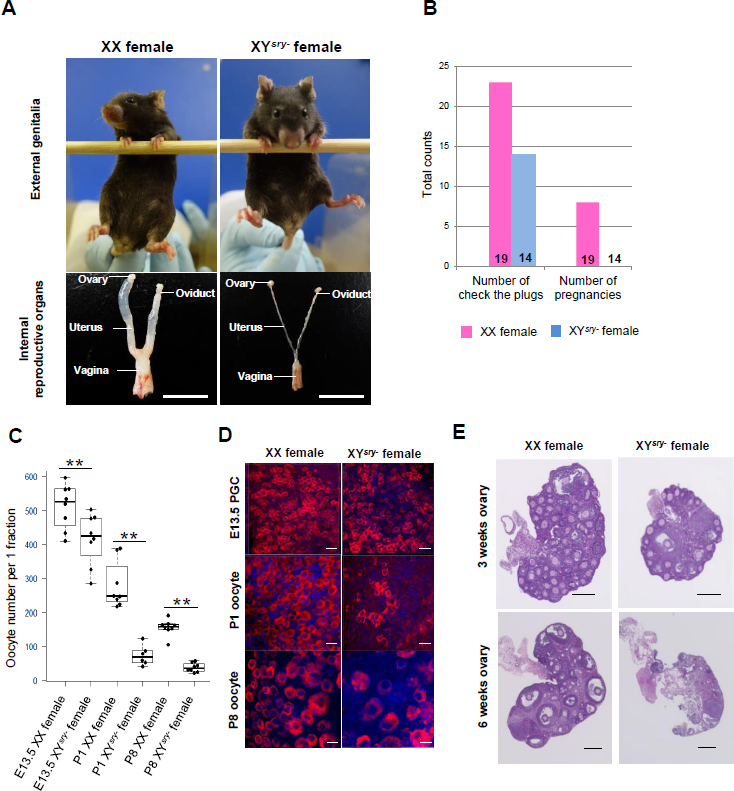
XY^*sry*−^-female mice exhibit infertility and oocyte depletion. (A) External genitalia and internal reproductive organs of an XX wild-type (WT) female (left) and an XY^*sry−*^-female (right) at 8 weeks of age. Generated XY^*sry−*^-female mice were characterised by a decreased anogenital distance and internal reproductive structure, comprised of a converted ovary. (B) Fertility test of XX WT and XY^*sry−*^-females. Left bar graph shows the total number of checked plugs; right bar graph illustrates the total number of pregnancies in each genotype. Number of tested females of each genotype is shown at the bottom of the respective bars. Pink and blue bars indicate WT and XY^*sry−*^-females, respectively. (C) Boxplot of the total oocyte number per 1 fraction (x: 212.2 μm × y: 212.2 μm × z: 25.0 μm) in XX WT and XY^*sry*−^-female gonads (Student’s *t*-test: *\*\*P* < 0.01 for comparisons of the germ cell numbers made at E13.5, P1, and P8). (D) Representative images of immunostained germ cells at different developmental stages (E13.5, P1, and P8) in XX WT and XY^*sry*−^-female gonads. PGCs and oocytes were stained with an antibody against a germ cell-specific marker (DDX4) and counterstained with DAPI (blue). Scale bar, 20 μm; magnification, ×40. (E) Representative haematoxylin and eosin-stained sections of ovaries of 3- and 6-week-old XX WT and XY^*sry−*^-females are shown. Scale bar, 200 μm.

### Transcriptome analysis of XY^*sry−*^-germ cells

To understand the mechanisms underlying the depletion of germ cells in XY^*sry−*^-females, we conducted transcriptome analysis followed by bioinformatics analysis. The RNA-Seq datasets provided accurate gene expression profiles: a summary of the gene expression profiles is shown in Table S2 and S3. Hierarchical cluster analysis revealed that the gene expression profiles segregated into two groups representing E13.5 PGCs and P1 oocytes and that both profiles of the XY^*sry−*^-female germ cells could be clearly distinguished from those of XX females and XY males (Fig. 2A, Fig. S2). Notably, the transcriptome data showed that many of the female and male PGC-specific expressed genes (FSGs, n = 245/651) and (MSGs, n = 157/428), which were identified by a previous study (Sakashita et al., 2015), were significantly up- and down-regulated, respectively, in XY female PGCs (Fig. S3, Table S4). Therefore, it is considered that XY female PGCs, contrary to its genetic sex, have been already switched at this stage from the male property to the female property.

**Fig. 2.**
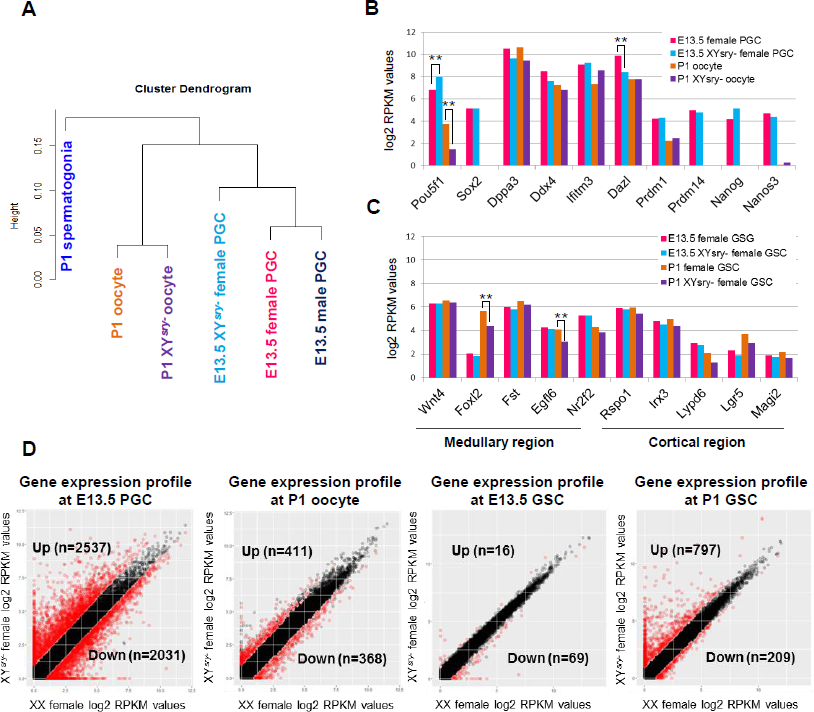
Extensive transcriptional changes in X XY^*sry*−^-female germ cells. (A) Unsupervised hierarchical cluster of all transcript profiles (*n* = 36,172) from 6 RNA-Seq datasets. The height of the vertical axis indicates the Euclidean distance (correlation distance) between objects. (B) Germline-specific gene expression patterns in each germ cell RNA-Seq dataset. Asterisks indicate statistically significant differences between XX WT and XY^*sry−*^-females (moderated *t*-test: \*\**P* < 0.01). (C) Gonadal somatic cell (GSC) marker gene expression patterns (5 medullary cell markers and 5 cortical cell markers) in each GSC RNA-Seq dataset. Asterisks indicate statistically significant differences between WT and XY^*sry−*^-females (moderated *t*-test: \*\**P* < 0.01). (D) The left two panels illustrate scatter plots of log2 RPKM values of WT (X-axis) and XY^*sry−*^-germ cell RNA-Seq samples (Y-axis) obtained at E13.5 and P1. The right two panels illustrate scatter plots of log2 RPKM values of WT (X-axis) and XY^*sry−*^-GSC RNA-Seq samples (Y-axis) obtained at E13.5 and P1. Red circles indicate genes found to be differentially expressed (≥ two fold difference in the expression level) according to the moderated *t*-test at *P* < 0.05.

The expression of marker genes for PGCs (Fig. 2B) and gonadal somatic cells (Fig. 2C) showed no global or significant change. However, the expression of POU domain, class 5, transcription factor 1 (*Pou5f1*), a transcription factor necessary for germ cell differentiation and survival, was significantly up- and down-regulated in E13.5 PGCs and P1 oocytes of XY^*sry−*^-females, respectively, and that deleted in azoospermia-like (*Dazl*), a RNA binding protein and critical for pluripotency maintenance, was significantly decreased in XY^*sry−*^-PGCs (Fig. 2B). Furthermore, the expression of forkhead box L2 (*Foxl2*), a gene encoding a forkhead transcription factor, and that of EGF-like-domain, multiple 6 (*Egfl2*), encoding a protein with a suggested an adhesive function, from medullary region of the ovary was significantly decreased in P1 XY^*sry−*^-female gonadal somatic cells (GSCs, Fig 2C). These findings demonstrate that the transcriptome datasets obtained were of high quality and valid for subsequent analysis.

Next we focused on differentially expressed genes in E13.5 XY^*sry−*^-female PGCs, which was screened based on 2 parameters, namely a fold change (FC) of >2 and the statistical significance determined using a moderated *t*-test with a Benjamini-Hochberg FDR of <0.05 (Fig 2D). In E13.5 XY^*sry−*^-female PGCs, 2,537 (21% of all transcripts) and 2,031 (17% of all transcripts) genes were up-and down-regulated, respectively (Fig 2D and Table S4). While, the number of genes screened as up- and down-regulated was decreased to 411 (3.2%) and 368 (2.9%) at the P1 oocytes (Fig 2D and Table S5). Most of the genes, whose expression was significantly changed, were classified to protein-coding cluster (84%-98% of all transcripts). Interestingly, proportion of non-coding RNAs (11.3%) was apparently high in the down-regulated genes of E13.5 XY^*sry−*^-female PGCs (Fig S3A). The 20 genes, which showed the greatest statistical significance in the PGCs and oocytes in XY^*sry−*^-females, are also shown in Fig S3B and C. These are reference gene types but the function in germ cells is not well known. Top 3 up-regulated genes of P1 XY^*sry−*^-female oocytes were expressed from Y chromosome (Fig S3). A few number of differentially expressed genes (16 up-regulated and 69 down-regulated) were detected in gonadal somatic cell at E13.5 XY^*sry−*^-female PGCs (Fig 2D and Table S6). However, the number of the genes greatly increased at P1 stage, especially in up-regulated genes (797 up-regulated and 209 down-regulated, Fig 2D and Table S7). This suggests functional defects of the survived P1 oocytes. These results demonstrated that sex-reversed germ cells are concealed serious transcriptional defects.

Significant changes of epigenetic modification related genes would affect a large scale change of gene expressions. Then, we examined expressions of DNA methylation and histone medication related genes in E13.5 XY^*sry−*^-female PGCs. No significant change was found in DNA methylation and demethylation related genes, except for *Dnmt3b* and *Tet3,* whose expression was very low even in WT female PGCs. However, interestingly, expression of genes, which are related to both active and negative markers of histone modifications, was significantly changed in many cases (Fig. S4). Importantly, expression of *Suv39h1,* H3K9 methylase, and *Kdm4a,* H3K9 demethylase, showed an opposite direction change in XY^*sry−*^-female, which would accelerate H3K9 demethylation of the germ cells. In addition, *Kdm6a,* H3K27 demethylase was up-regulated and *Ring1a,* H2AK119 ubiquitin ligase, was down regulated.

### Cause of PGC and oocyte depletion

The number of PGCs at E13.5, when they had reached their maximum (Yokobayashi et al., 2013), was already significantly fewer in XY^*sry−*^-females (Fig. 1C). This observation suggested that the PGCs exhibited inferior proliferation compared with that of XX females, and also that cell death program was accelerated. Furthermore, the number decreased to 1/5 at the P1 stage in XY^*sry−*^-females compared with XX females. Subsequent analysis of gene ontology (GO) and functional pathways using differentially expressed gene sets provided valuable information toward understanding the mechanisms underlying germ cell depletion in XY^*sry−*^-females (Table S8 and Figure S5). Up-and down-regulated genes were respectively enriched for the following biological processes (Fig. S5A-B): up-regulated genes; “Tissue morphogenesis” and “Enzyme linked receptor protein”, down-regulated genes: “DNA metabolism”, “Cell cycle process” and “Chromosome organization”. GO term interaction analysis formed three large networks regarding “Regulation of gene expression”, “Tissue morphogenesis” and “Cell cycle and Cell division” (Fig. S5C).

In particular, the many “Wnt/β-catenin signal”-related genes were up-regulated (Fig 3A) in PGCs of XY^*sry−*^-females at E13.5. Furthermore, the expression of β-catenin (*Ctnnb1*) was not changed (Table S2); however, immunofluorescence staining revealed that the proteins were specifically located in the nuclei of PGCs in XY^*sry−*^-females, but not in XX-females (Fig 3B). This observation suggests an adverse effect of β-catenin on cell proliferation. Conversely, the result that many genes related to meiosis were down-regulated suggested that the transition of PGCs to meiosis at E13.5 was delayed in XY^*sry−*^-females (Fig 3C). In addition, Immunofluorescence staining showed that many PGCs in XY^*sry−*^-females expressed the mitosis marker, *Ki67* (Fig 3D). Furthermore, apoptosis-related genes, especially the *Tnf* family, were enhanced (Fig S6A) with tumor necrosis factor receptor superfamily member 12A (*Tnfrs12a*), exhibiting a high degree of elevated expression. Additionally, mRNA expression of the cell death makers *Traf, Ripk2*and *Caspase* tended to increase in XY^*sry−*^-female PGCs as well. However, although the apoptotic cell populations were slightly increased in E13.5 XY^*sry−*^-PGCs, these were not statistically significant compared with that of WT controls (Figure S6B-C). Unnaturally, many endogenous retrovirus (ERVs) and LINE1 retrotransposons were unlikely highly expressed in E13.5 XY^*sry−*^-female PGCs (Fig. 3E). ERV1 and satellite DNA were up-regulated in P1 oocytes. Furthermore, α-thalassemia mental retardation X-linked (*Atrx*), and suppressor of variegation 3-9 homolog 1 (*Suv39h1*), which are involved in retrotransposable element silencing and chromatin remodelling of heterochromatin structure, were markedly repressed in E13.5 XY^*sry−*^-PGCs (Fig. 3F). These results suggested that oocyte cyst breakdown and primordial follicle formation were considerably disrupted in the neonatal XY^*sry−*^-ovary. Then, we asked whether this aberrant expression of retrotransposable elements led to meiotic defects in the foetal germ cells of XY^*sry−*^-females. Immunofluorescence staining of synaptonemal complex protein 3 (SCP3) showed that the synaptonemal complex between homologous chromosomes was clearly observed in both XX and XY^*sry−*^-germ cells at E17.5 (Fig. 3G). However, fluorescent signals of phosphorylation of histone H2AX (γH2AX) foci, which indicates unpaired regions, were detected only in the XY^*sry−*^-germ cells (25/25; 100%). Furthermore, SCP3 and synaptonemal complex protein 1 (SCP1) were well co-localized across a wide range of chromosomes in XX germ cells; in contrast, the co-localization was observed in only a few cases (6/24; 24%) in XY^*sry−*^-germ cells. Thus, the meiotic process in XY^*sry−*^-germ cells appeared to be disturbed, accompanied with chromosomal asynapsis.

**Fig. 3.**
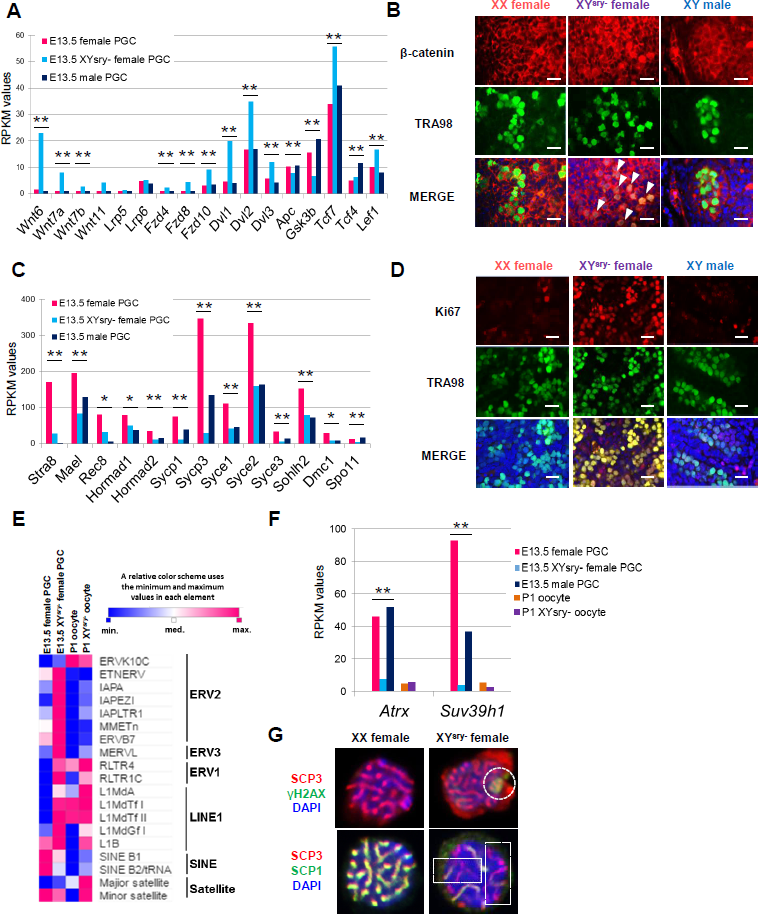
Meiotic defect of XY^*sry−*^-female germ cells accompanying impairment of cell cycle progression and retrotransposable element activation. (A) Expression patterns of Wnt signalling pathway components in each E13.5 PGC RNA-Seq dataset. Asterisks indicate statistically significant differences between RPKM values in E13.5 female, XY^*sry−*^-female and male PGCs (one-way ANOVA, \*\**P* < 0.01). (B) Representative images of immunostained E13.5 gonads. PGCs were stained with antibodies against a germ cell-specific marker (TRA98) and α-catenin, and counterstained with DAPI (blue). Arrowheads show XY^*sry−*^-female PGCs expressing stabilised β-catenin in the nucleus. Scale bar, 20 μm; magnification, ×40. (C) Expression patterns of meiosis-related genes in each E13.5 PGC RNA-Seq dataset. Asterisks indicate statistically significant differences between RPKM values in E13.5 female, XY^*sry*−^-female and male PGCs (one-way ANOVA: \**P* < 0.05, \*\**P* <0.01). (D) Representative images of immunostained E14.5 gonads. PGCs were stained with anti-TRA98 and anti-Ki67 (cell proliferation marker, red) antibodies, and counterstained with DAPI. Scale bar, 20 μm; magnification, ×40. (E) Expression of representative retrotransposable elements in E13.5 and P1 germ cells. Data are represented by a heat map of the relative expression levels of each retrotransposable element. (F) Expression of *Suv39h1* and *Atrx* in each germ cell RNA-Seq dataset. Asterisks indicate statistically significant differences between RPKM values in E13.5 female, XY^*sry−*^-female and male PGCs (one-way ANOVA, \**P* < 0.01). (G) Representative images of immunostained E17.5 pachytene stage oocytes from WT (left) and XY^*sry−*^-(right) females. Top panel: oocytes were stained with anti-SCP3 (red) and anti-γH2AX (green) antibodies, and counterstained with DAPI. Dashed circle indicates yH2AX-positive region. No yH2AX signal was detected in XX WT oocytes (0/25; 0%), whereas all XY^*sry−*^-oocytes were positive for yH2AX (25/25; 100%). Bottom panel: Oocytes were stained with anti-SCP3 (red) and anti-SCP1 (green) antibodies and counterstained with DAPI. Dashed rectangles indicate the asynaptic regions. In XX WT oocytes, all homologous chromosomes have synapsed completely along their length at the pachytene stage. However, approximately a quarter of XY^*sry−*^-oocytes exhibited extensive chromosomal asynapsis (6/25; 24.0%).

Notably, histological analysis (Fig. 4A) and immunofluorescence staining for DDX4 (Fig. 4B) demonstrated that only a few oocytes were observed in the medullary region of P1 XY^*sry−*^-ovary. MT staining showed that the connective tissues and collagen fibers were proliferated in the medullary region (Fig. 4A). Notably, XY^*sry−*^-oocytes in the cyst were apparently fewer than those of WT (Fig. 4A) and the proportion of the oocytes in the cyst was significantly high (Fig. 4C). Furthermore, a significantly high proportion of apoptotic oocytes was detected in the cortical region of control XX ovary but only few in the XY^*sry−*^-ovary (Fig 4B, D, Fig. S7). These observations were supported by transcriptome data. Many folliculogenesis-related genes were down-regulated in P1 XY^*sry−*^-oocytes (Fig. 4E). GO analysis showed that differentially expressed genes in the P1 oocytes were enriched in particular annotations, such as “Cell adhesion” and “Cell cycle” (Table S8). Histopathological analysis demonstrated that the primordial oocyte reserve population had almost disappeared from the medulla of the P8 XY^*sry−*^-ovary (Fig. 4E-F). However, the primary follicles were localized within the cortical region differ from the XX female ovary.

**Fig. 4.**
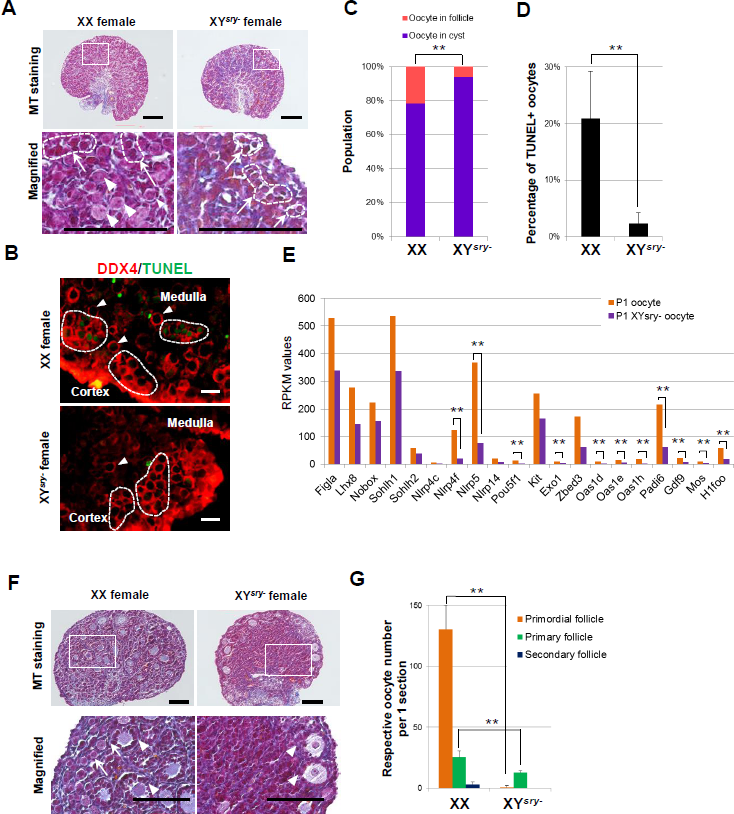
Retardation of oocyte cyst breakdown and elimination of primordial follicles in neonatal XY^*sry−*^-ovaries. (A) Representative Masson Trichrome (MT) sections of ovaries of P1 XX WT and XY^*sry−*^-females are shown. Bottom panels are magnified images of dashed rectangle depicted in upper panels. Arrows/dashed circles and arrowheads indicate oocyte cysts and primordial follicles, respectively. Scale bar, 100 μm. (B) TUNEL assay. Representative images of immunostained P1 ovaries in XX WT and XY^*sry−*^-females. Ovaries were stained with antibody against a germ cell-specific marker (DDX4, red) and TUNEL (apoptotic cell marker, green). Dashed circles indicated by an allow indicate oocyte cysts and arrowheads indicate primordial follicles. Scale bar, 20 μm; magnification, ×40. (C) Percentage contribution of oocyte in cysts and oocyte in follicles from P1 XX and XY^*sry−*^-ovaries. Asterisk indicates statistically significant differences between the groups in the rating of “oocyte in cysts” and “oocyte in follicles (Chi-squared test: \*\**P* < 0.01, n = 3). (D) Percentage of TUNEL positive oocytes in P1 XX and XY^*sry−*^-ovarian section. The percentage of apoptotic oocytes (TUNEL positive) was shown as the mean ± standard error of the mean. Asterisk indicates a statistically significant difference (Student’s *t*-test: \*\**P* < 0.01, *n* =3). (E) Expression patterns of folliclogenesis-related genes in each P1 oocyte RNA-Seq dataset. Asterisks indicate statistically significant differences between RPKM values in XX WT and XY^*sry−*^-oocytes (moderated *t*-test: \*\**P* < 0.01). (F) Representative MT sections of ovaries of P8 XX WT and XY^*sry−*^-females are shown. Bottom panels are magnified images of the dashed rectangle depicted in the upper panels. Arrows and arrowheads indicate primordial follicles and primary follicles, respectively. Scale bar, 100 μm. (G) Number of respective oocytes per 1 ovarian section at P8. Asterisks indicate a statistically significant difference between XX WT and XY^*sry−*^-oocytes (Student’s *t*-test: \*\**P* < 0.01, *n* = 3).

### Y-chromosome-linked gene expressions

Considering the fertility of XO mice, it may be considered likely that the dysfunction of *Sry* deficient XY^*sry−*^-oocytes is caused by the existence of the Y chromosome itself; i.e., that the expression of genes located on the Y chromosome might present a key factor. Single-copy genes located on Y chromosome MSYp region were expressed at similar levels to those of XY male germ cells, except some cases (Fig 5B). For example, the expression of *Zfy1, Elf2s3y* and *Uty* was significantly low in E13.5 XY^*sry−*^-PGCs. The multiple copy genes *Rbmy* also showed significantly decreased expression in XY^*sry−*^-germ cells, especially at P1. These results suggest that the expression of particular genes from the Y chromosome in XY^*sry−*^-females represented a trigger for the depletion and dysfunction of XY^*sry−*^-female germ cells.

**Fig. 5.**
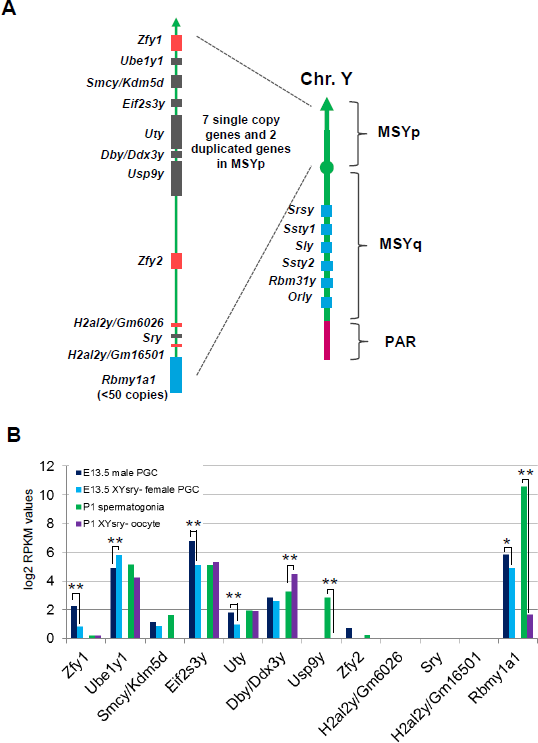
Expression of Y-linked genes in E13.5 PGCs and P1 oocytes of XY^*sry−*^-females. (A) Schematic diagram of the mouse Y chromosome showing the male-specific region of the Y short arm (MSYp; represented to scale in the magnified view), male specific region of the Y long arm (MSYq), and pseudoautosomal region (PAR). The MSYp region has seven single copy genes, two duplicated genes, and one multi-copy gene. The MSYq region carries several multi-copy genes. The PAR is the chromosomal region, where sex (X-Y) chromosome pairing occurs. (B) Expression patterns of Y-linked genes from E13.5 and P1 germ cell RNA-Seq datasets.

### Mature oocyte dysfunction

XY^*sry−*^-females were infertile but possessed the antral follicle oocytes at 3-weeks old (Fig. 1E). Therefore, we first examined whether the mutants ovulate mature oocytes in response to exogenous gonadotropins at 4 weeks old. The number of ovulated XY^*sry−*^-oocytes was less than half of the WT (XX: 51.5 ± 2.5 vs XY^*sry−*^: 20.5 ± 9.1, Student’s *t*-test *P* < 0.01). The MII oocytes exhibited aberrant spindle formation: diffused poles, skewed spindles, disrupted spindles and misaligned and decondensed chromosomes (Fig. S8B-C). After *in vitro* fertilization (n = 3), 35.6% (16/32) of the MII oocytes showed the capability to fertilize and form male and female pronuclei (Fig. S8D). Of these 10 were arrested at 2-cell stage (Fig. S8D), and one blastocyst obtained but Immunostaining with anti-Oct4 showed failure of differentiation into inner cell mass and trophectoderm cells (Fig. S8E).

To unravel the molecular mechanisms underlying the mature oocyte dysfunction in XY^*sry−*^-females, we conducted a single-oocyte transcriptome analysis using 27 XY^*sry*−^” and 30 XX oocytes. Principal component analysis (PCA) revealed an almost identical gene expression profile in XX single oocyte libraries (Fig. 6A). However, each XY^*sry−*^-oocyte profile was largely varied in the second principal component (PC2) (Fig. 6A). The maternal effect genes, which are important for early development, were expressed at levels similar to those of XX oocytes (Fig. 6B). Clustering analysis revealed that 15 XY^*sry−*^-oocytes were classified into a cluster that was clearly distinguished from the WT cluster consisting of the 29 oocytes (Fig. 6C). The other mutant oocytes formed a sub-cluster differing from the WT cluster. GO analysis using differentially expressed gene sets in XY^*sry−*^-oocyte, provided the terms related to the oocyte dysfunctions (Fig. 6C); for example, for up-regulated genes: “meiotic cell cycle” and “apoptotic process”, down-regulated genes: “protein ubiquitination”, “sister chromatid cohesion” and “meiotic nuclear division”. Furthermore, interestingly, the retrotransposable elements such as LINE1, ERV and satellite DNA were highly expressed in 18 XY^*sry−*^-oocytes, whereas only a single WT oocyte was classified in this cluster (Fig. 6D, Fisher’s exact test, *P* < 0.01). Thus, single oocyte RNA-Seq data demonstrated that most of XY^*sry−*^-MII oocytes exhibited extensive transcriptional anomalies, which might responsible for the observed impairments in the progression of second meiotic division and early development.

**Fig. 6.**
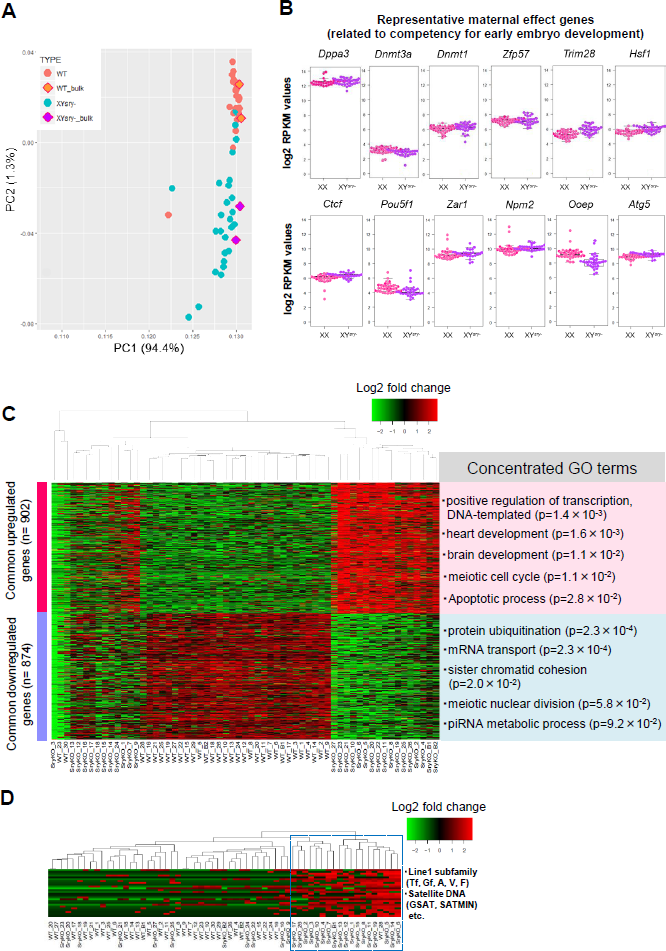
Gene expression profiling of MII oocytes from XY^*sry*−^-females by single cell transcriptional analysis. (A) Bi-dimensional PCA of gene expression profiles of XX WT (n = 30) and XY^*sry*−^-(n = 27) single oocyte. The bulk oocytes (20 oocytes, n = 2) data were shown in square. The first principal component (PC1) captures 94.4% of the gene expression variability and the second principal component (PC2) captures 1.3% oocytes, respectively. (B) Boxplots of log2 RPKM values for 12 representative maternal effect genes in XX WT (pink) and XY^*sry−*^-oocytes (purple). No significant difference was observed for each gene expression. (C) Heatmap shows relative expression of the 902 commonly up-regulated and 874 down-regulated genes in each XY^*sry−*^-MII oocyte identified by k-means clustering. Representative GO terms and the enrichment P values are shown at right. (D) Heatmap shows relative expression of the 20 L1 subfamilies, 3 ERV2 elements and 3 satellite DNAs identified using k-means clustering methods, in XY^*sry−*^-MII oocytes. Blue dashed rectangle indicates a cluster, consisting of highly expressed retorotransposable elements. Note, XY^*sry−*^-MII oocytes were highly assigned to this cluster (18/19 (94.7%), Fisher’s exact test; *P* < 0.01).

### Discussion

Germ cells from any type of sex-reversed mice, i.e., XX males and XY females, exhibit morphologically normal sexual features; however their germ cell development results in failure and inevitably they are infertile (Kato et al., 2013; Lovell-Badge and Robertson, 1990; Ma et al., 2000; Mahadevaiah et al., 1993; Taketo, 2015; Vanderhyden et al., 1997; Vernet et al., 2012; Vernet et al., 2011; Vernet et al., 2014b). Here, we present the first demonstration of the molecular mechanisms underlying germ cell depletion and dysfunction in XY^*sry−*^-female mice. In female mice, PGC precursors become committed to differentiate down a female pathway into oocytes between E12.5 and E13.5 (Suzuki et al., 2015), which is typically accompanied by a transition from mitosis to meiosis. From the present integrated analysis of germ cells from XY sex-reversed females, the complicated mechanisms underlying the observed oocyte depletion in the ovary may be explained by two major distinguishable processes; a decrease of PGCs following the transition from mitosis to meiosis, and a rapid loss of the primordial oocyte pool after birth.

For the first process, it is considered that the Wnt/β-catenin signalling pathway would likely be involved in the decline of PGC proliferation in the XY^*sry−*^-females as a large number of genes involving this pathway exhibited significant enhancement. However, although the expression of β-catenin itself was not increased, it has been reported that the stable localization of β-catenin at a nucleus of mitotic stage PGCs blocks the transition from G1 to S stage and leads to inhibitory effects on germ cell proliferation in mice (Kimura et al., 2006). Consistent with this observation, fluorescence microscopy clearly showed the predominant nuclear localization of β-catenin in the XY^*sry−*^-female PGCs. Furthermore, the nuclear localization of β-catenin is inhibited by phosphorylation of exon 3 of this protein, a function catalysed by glycogen synthase kinase 3β (GSK-3b) (Wu and Pan, 2010). Therefore, the finding that *Gsk3b* expression was decreased in XY^*sry−*^-female PGCs suggested that β-catenin localization into the nuclei of these cells was facilitated likely extensively impacting gene regulation. In tumor cells, overexpression of β-catenin induce a delay in the cell cycle and apoptosis, independent from the actions of *TCF* and *LEF1* actions (Kim et al., 2000; Olmeda et al., 2003). Therefore, we considered that the Wnt/β-catenin signal pathway represented a critical factor towards decreasing the proliferation of PGCs in XY^*sry−*^-females.

In mice, it is known that after PGCs reach to the maximum number (12,000) and enter the prophase of meiosis I at E13.5-14.5, they then enter a continuously decreasing stage resulting in less than one third of the maximum number of cells remaining around the time of birth (Lei and Spradling, 2016; Pepling and Spradling, 2001; Yokobayashi et al., 2013). Evidence has shown that apoptosis represents a major cause of this germ cell reduction. Our previous transcriptome study in mice demonstrated that an intrinsic pathway, such as the mitochondrial pathway, is required to induce the apoptosis of female PGCs; however, the extrinsic pathway was not activated (Sakashita et al., 2015). We also found that the mitochondrial pathway of apoptosis was activated in the PGCs of XY^*sry−*^-females as well. Notably, the present results further demonstrated that in addition to the mitochondrial pathway, an extrinsic apoptosis pathway was also activated in XY^*sry−*^-female PGCs, in which the expression of *Tnf* receptors was significantly enhanced. This additional pathway may therefore promote apoptosis and result in the loss of greater number of oocytes. Furthermore, the enrichment of differentially expressed genes among the ‘Response DNA damage stimulus’ and ‘DNA repair’ categories might also have been involved in the germ cell loss. In particular, the down-regulation of *Rad51* and *Brca,* which encode DNA repair enzymes (Scully et al., 1997), in XY^*sry−*^-female PGCs would be expected to induce apoptosis as well. For example, it has been reported that cells mutated for the BRCA proteins lack DNA repairing ability (Yoshida and Miki, 2004), and that *Rad51* is involved in the resistance to apoptosis following DNA damage, and promotes homologous recombination by chromatid exchange mechanism (Arnaudeau et al., 1999; Xia et al., 1997).

In comparison, in XO female mice, it has been shown that the oocyte number reduced to nearly half compared with that of XX females at neonatal and developmental periods (Burgoyne and Baker, 1981, 1985). Recently, Cloutier et al. (2015) showed that the oocyte elimination in XO mice is linked to the presence of H2AFX serine-139 phosphorylation on the chromatin of asynapsed chromosomes (Cloutier et al., 2015). Therefore, the existence of asynapsed XY chromosomes may be related to oocyte elimination in sex-reversed females as well. Consistent with this suggestion, we detected γH2AX foci in all pachytene oocytes from E17.5 XY^*sry−*^-females. In male spermatogenesis, germ cells that fail to synapse their chromosomes or fail to undergo meiotic sex chromosome inactivation are eliminated by apoptosis during the mid-pachytene stage (Burgoyne et al., 2009; Vernet et al., 2011), and these phenomena were regulated by *Zfy2,* which is Y-linked transcription regulator (Vernet et al., 2011). This suggests that *Zfy2* may be involved in asynapsed pachytene oocyte depletion in XY^*sry−*^-females. Moreover, we demonstrated a marked increase of LINE-1 and ERV retrotransposons (>10 fold) in XY^*sry−*^-females, which might have accelerated the germ cell depletion. For example, it has been shown that the forced enhancement of L1 expression is involved in foetal oocyte attrition through increasing the nuclear level of LINE1, which induces a failure of meiotic progression (Malki et al., 2014).

Oocyte loss in the XY^*sry−*^-females observed in postnatal stage was drastic and distinctive; thus, the oocyte cyst breakdown may be closely related. It is known that *Foxl2* positive medullary granulosa cells are important for precise oocyte cyst breakdown and primordial follicle formation (Tingen et al., 2009; Wang et al., 2017). Knockout of *Foxl2* in granulosa cells impairs cyst breakdown by inhibiting granulosa differentiation and proper laying down of the basal lamina around the forming follicle (Uda et al., 2004). In this study, the expression of *Foxl2* was significantly decreased in XY^*sry−*^-female GSCs, compared with WT (Fig. 2C). This might be related to excess proliferated connective tissues and collagen fibres that filled the medullary region of the P1 XY^*sry−*^-ovary (Fig. 4A). Thus, regression of granulosa cells and the decrease of *Foxl2* expression might be involved in impairment of oocyte cyst breakdown and primordial follicle formation in the neonatal XY^*sry−*^-ovary. Furthermore, it was also reported that *Foxl2* deficiency is also associated with premature ovarian failure (Crisponi et al., 2001; Uda et al., 2004). From these, it is supposed that the follicles distributed into the cortical region were aberrantly activated by decreased expression of *Foxl2,* resulting in the precocious exhaustion of XY^*sry−*^-oocytes prior to the pubertal stage.

The fertility of XO female mice strongly indicates that genes expressed from the Y chromosome are involved in oocyte depletion in XY^*sry−*^-females. However, little is known regarding the function of these genes other than *Sry.* To date, mice harbouring many types of Y chromosome mutation have been reported (Royo et al., 2010; Vernet et al., 2011; Vernet et al., 2014b). Of these, the infertility of XO mice expressing *Zfy2* indicates that *Zfy2* can serve as a restricted factor if it is expressed in oocytes (unlike the situation observed in our XY sex-reversed females) (Vernet et al., 2014b). Sex reversed XX male mice harbouring *Sry* transgene resulted in male infertility consequent to a failure of spermatogenesis (Koopman et al., 1991). However, X^E^OSry mice carrying the X-linked *Eif2s3y* gene occasionally led to formation of haploid spermatids, which could be used for round spermatid injection (Vernet et al., 2011; Yamauchi et al., 2014). In addition, Lavery et al. (2011) reported that *Sox9* deletion resulted in sex-reversed XY females that exhibited subfertility in some cases (Lavery et al., 2011), although a comparison between these mice and XY^*sry−*^-females has not been concluded. These findings suggest that Y-linked genes might cause detrimental effects on oogenesis. Accordingly, our transcriptome analysis revealed that the *Ube1y1, Eif2s3y, Uty,* and *Ddx3y* genes expressed in XY male PGCs were also expressed at similar levels in XY^*sry−*^-female PGCs (Fig. 5B). From these findings, it might also indicate that if particular Y-linked genes were deleted in addition to Sry, fertile germ cells might be obtained from sex reversed XY females.

Furthermore, our single cell transcriptome analysis provided evidence to understand the molecular mechanisms underlying the dysfunction of XY^*sry−*^-oocytes. Natably, developmental process-related genes, which were likely to remain transcriptionally silent in germline cells through bivalent chromatin marks with H3K4me3 and H3K27me3 (Hammoud et al., 2009; Reik, 2007; Sachs et al., 2013), were activated in XY^*sry−*^-MII oocytes (Fig. 6C). Moreover, 6 genes (Zfy1, *Ube1y1, Kdm5d, Uty, Ddx3y, Zfy2*) located at MSYp region also expressed in XY^*sry−*^-mature MII oocytes. As a result of GO analysis for respective using the DAVID web tool (Wang et al., 2013), it was revealed that *Ddx3y* and *Eif2s3y* had the functions associated with “chromosome segregation” and “translation”, respectively. Furthermore, it had been reported that the expression of *Zfy2* in XY oocytes elicited frequent MII defects and preimplantation developmental arrest.

In conclusion, the present study conducted inclusive transcriptome and bioinformatics analysis and demonstrated that the serious germ cell loss in XY sex-reversed female mice is induced by a decline of cell proliferation, an acceleration of cell death programs and follicle formation. Furthermore, we showed that the failure of DNA remodeling and repair in XY^*sry−*^-female oocytes resulted in germ cell loss and dysfunction. These results may provide further insight into understanding disorders of sex development in human, such as Swyer syndrome, wherein the patients are anatomically females but have an XY karyotype and exhibited infertility consequent to mutations in *Sry* and *Map3k1* (Pearlman et al., 2010).

## Materials and Methods

### Animals and ethic statements

This study was carried out in strict accordance with the Tokyo University of Agriculture Guide for Care and Use of Laboratory Animals. The protocol was approved by the Committee on the Ethics of Animal Experiments of the Tokyo University of Agriculture (Permit Number: 260064SE). At the time of sample collection, all animals (CLEA JAPAN, JP) were sacrificed by cervical dislocation, and all efforts were made to minimize suffering.

### Generation of *Sry* mutant mice (XY^*sry−*^-females) by CRISPR/Cas9 system

To generate the *Sry* targeting vector, we used a pX330-U6-Chimeric BB-CBh-hSpCas9 (pX330) plasmid, kindly provided by Dr. Ikawa (Research Institute for Microbial Diseases, Osaka University). sgRNA sequence was designated on SRY HMG-box domain (5’– TGGTGTGGTCCCGTGGTGAG –3’), and inserted annealed double strand DNA with 4 overhangs into BbsI site in pX330 plasmid. B6 female mice were superovulated and mated B6 male mice, and zygotes were collected from ampulla of the oviducts. The microinjection of the *Sry* targeting pX330 plasmid (2 ng/μl) into male pronuclear of PN3-5 stage zygotes was carried out under standard procedures. The injected zygotes were cultured in potassium simplex optimized medium (KSOM) at 37°C and embryos developed into the two-cell stage were transferred into oviducts of pseudopregnant ICR female mice.

### PCR-based genotyping and sexing

Genomic DNA was extracted from heads or tail tips using NaOH extraction method (Wang et al., 1993). For genotyping of Sry, PCR was carried out using specific primer sets: Sry_F (5’– GTGACAATTGTCTAGAGAGCATGGAGG –3’) and Sry_R (5’ – TCCAGTCTTGCCTGTATGTGAT – 3’). The PCR products were purified by QIAquick PCR purification kit (QIAGEN, Hilden, FRG) and used as templates for sequencing. Sanger sequencing was done by ABI PRISM 3100 Genetic Analyzer (Applied Biosystems, CA, USA). For PCR-based sexing, X chromosome specific gene (*Xist*) and Y chromosome specific gene (*Zfy*) were amplified using following primer sets (Kay et al., 1994; Zuccotti and Monk, 1995):

*Xist*_F; 5’− AGGATAATCCTTCATTATCGCGC −3’,

*Xist*_R; 5’− AAACGAGCAAACATGGCTGGAG −3’

Zfy_F; 5’− GACTAGACATGTCTTAACATCTGTCC −3’

Zfy_R; 5’− CCTATTGCATGGACAGCAGCTTATG −3’

### Fertility and development test

XY^*sry−*^-females, 8-12 weeks old, were mated with fertile B6 males. The presence of the vaginal plug was checked every morning for four weeks. MII oocytes were collected by PMSG-hCG treatment from 4-week-old mice, and the ovulated oocytes were applied to IVF. The zygotes obtained were subjected to *in vitro* culture for 4 days in KSOM.

### Germ cells and gonadal somatic cells collection

To collect E13.5 PGCs, P1 oocytes and gonadal somatic cells (GSCs), gonads from 13.5 dpc and 1 dpp after embryo transferred were digested 1 mg/ml collagenase solution (Wako, Oosaka, JP) at 37°C for 40 min, followed by treatment with 0.25% trypsin-EDTA solution (0.53 mM; Sigma, MO, USA) at 37°C for 15 min. After adding foetal bovine serum (FBS), a single-cell suspension was obtained by gentle pipetting. Cells were then incubated in a 1:50 dilution of PE-conjugated anti-SSEA1 (for labeled E13.5 PGCs, 560142, BD Pharmingen, NJ, USA) or PE-conjugated anti-CD117 (for labeled P1 oocytes, 105807, BioLegend, CA, USA) antibody. PE positive cells (PGCs and oocytes) and negative cells (Gonadal somatic cells) were isolated and collected using a FACSAria II cell sorter (BD Bioscience, NJ, USA, Fig. S9). Ovulated MII oocytes were extracted from the oviducts of XX and XY^*sry−*^-female mice according to methods described in our previous report (Obata and Kono, 2002).

Considering the variability in the genotype of the XY^*sry−*^-females, when collecting PGCs, oocytes, gonadal somatic cells and MII oocytes, we supplied pooled gonadal cell suspension and MII oocytes from 4 and 5 XY^*sry−*^-female individuals at each time point. The *Sry* mutation types are listed below. Furthermore, we confirmed that all mutant lines contained a frameshift mutation within HMG-box coding region, by converting DNA nucleotide sequences to amino acid sequences, and thus were expected to cause *Sry* protein disruption.

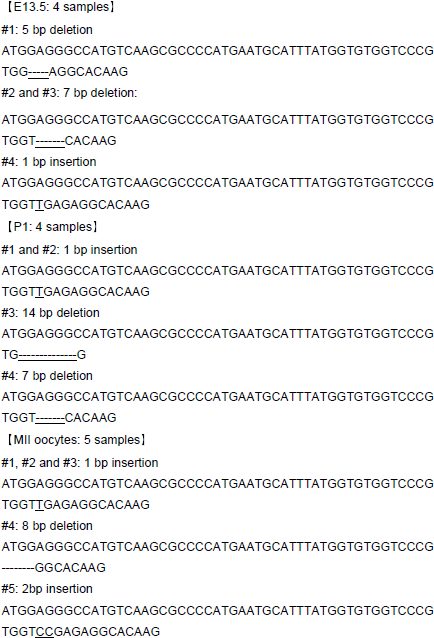

### RNA isolation, RNA-Seq library preparation and sequencing

Total RNA of PGCs, oocytes and GSCs was isolated using an RNeasy Micro Kit (QIAGEN, Hilden, FRG) with DNase treatment. cDNA synthesis and pre-amplification were performed with total RNA (10 ng) using a SMARTer Ultra Low Input RNA Kit and an Advantage 2 PCR Kit (Clontech, CA, USA), respectively, according to the manufacturers’ instructions. Pre-amplified cDNA was fragmented into 200-bp fragments using an S2 sonicator (Covaris, MA, USA) and then used to construct sequencing libraries using a NEBNext Ultra DNA Library Prep Kit, according to the manufacturers’ protocol (New England BioLabs, MA, USA). Two biological replicates were used for each sample.

To perform single cell RNA-Seq analysis of 4-weeks-old XX and XY^*sry−*^-MII oocytes, single and bulk (n = 20-22) MII oocytes from oviducts of the respective genotypes were picked by glass pipette and were immediately lysed in Clontech Lysis Buffer (Mountain View, CA, USA). The whole lysate was subjected to cDNA synthesis and RNA-Seq library using a SMARTer Ultra Low Input RNA Kit (Clontech) and Nextera XT library prep kit, according to manufacturer’s instruction (Illumina, San Diego, CA, USA). Indexed libraries were pooled (10 nM each), and sequenced using an Illumina Hiseq2500 sequencer (single-end, 100 bp condition).

### RNA-Seq alignments and statistical analysis

RNA-Seq reads for each sample were aligned to the mouse genome (mm10, Genome Reference Consortium Mouse Build 38) with the CLC Genomics Workbench (QIAGEN, Hilden, FRG). Aligned reads were subsequently assembled into transcripts guided by reference annotation (mm10, UCSC gene annotation). Transcript expression was quantified in terms of reads per million mapped reads and normalized using RPKM method with Strand NGS (Agilent, CA, USA). To identify significant differentially expressed genes between XX female vs XY^*sry−*^-female E13.5 PGCs and P1 oocytes, we used 2 selection criteria. The first criterion was a fold-change in expression of at least two-fold. The second criterion was removal of false positive genes using a moderated *t*-test with Benjamini-Hochberg false-discovery rate (FDR) of <0.05). Differentially expressed gene lists were used for GO analysis with the DAVID web tool (http://david.abcc.ncifcrf.gov/) (Huang da et al., 2009); a background of all mouse genes was applied. Biological-process term groups with a significance of *P* < 0.01 (modified Fisher’s exact test) were considered significant.

Single oocyte transcriptome datasets were applied to PCA and k-means cluster analysis to identify common up and downregulated genes in XY^*sry−*^-oocytes groups. Respective gene expression profiles were subjected to bi-dimensional PCA using R prcomp function and ggplot2 package (https://www.r-project.org/)

Each sample score from the covariance matrix was plotted in the two eigenvectors PC1 and PC2. To determine the optimum number of k-means clusters, we used the hierarchical cluster datasets of each gene, and further grouped the determined 4 k-means subclusters. The first subcluster comprised genes that showed common upregulated genes (n = 902) in XY^*sry−*^-MII oocytes. The second subcluster comprised genes showing common down-regulated genes (n = 874) in XY^*sry−*^-MII oocytes. In contrast, other two subclusters (third and fourth subcluster) comprised genes that showed uniformly expression pattern between XX and XY^*sry−*^-MII oocytes. We performed GO analysis of common up and down-regulated gens using the DAVID web tool.

### Transposable elements analysis

The current build of rodent repeat sequences was downloaded from Repbase (http://www.girinst.org/repbase/) and filtered for *Mus musculus* sequences. Raw RNA-Seq reads were aligned to the repetitive sequence database by CLC Genomics Workbench (QIAGEN, Hilden, FRG). Aligned reads were subsequently assembled into transcripts guided by reference annotation. Transcript expression was quantified in terms of reads per million mapped reads and normalized using RPKM method with Strand NGS (Agilent, CA, USA).

### Tissue collection, histological analysis and TUNEL assay

We fixed freshly dissected ovaries in 4% paraformaldehyde in PBS (−) at room temperature for overnight. The following day, fixed tissue samples were subjected to paraffin sectioning, hematoxylin and eosin (HE) staining and Masson’s trichrime (MT) staining. Also, TUNEL positive oocytes were detected in paraffin sections of P1 ovaries using the anti-DDX4 antibody (1/300 dilution, ab13840, Abcam, Cambridge, UK) and In Situ Cell Death Detection Kit, Fluorescein (11684795910, Roche, Basel, Switzerland).

### Germ cell counts

The number of germ cells was determined for each day of development from 13.5 dpc, 1 dpp and 8 dpp. First, to visualize germ cell, each whole ovary was stained sequentially first with anti-DDX4 antibody (ab13840, Abcam, Cambridge, UK) then the anti-rabbit Alexa 568 secondary antibody. Second, the number of DDX4 positive cells in a single representative section was counted. Total number of germ cells in one fraction was calculated using stacks of confocal sections (LSM710, Zeiss, Oberkohen, FRG).

### Immunofluorescence staining

Whole gonads were fixed in 4% paraformaldehyde in PBS (−) for 1 hr, and then incubated for 2 days at 4 °C with blocking solution (10% FBS, 3% BSA, 0.2% TritonX-100 in PBS (−)). Whole mount immunofluorescence analysis conducted using the following primary antibodies: anti-β-catenin (1/200 dilution, 610153, BD Transduction Laboratories, NJ, USA), anti-Ki67 (1/200 dilution, 556003, BD Pharmingen, NJ, USA), anti-cCASP3 (1/200 dilution, 9661, CST, MA, USA), anti-cPARP (1/200 dilution, 9544, CST, MA, USA) and anti-TRA98 (1/200 dilution, 73-003, BioAcademia, Oosaka, Japan). Gonads were incubated for 3 days at 4 °C with primary antibody, and then washed and incubated with Alexa Fluor 488- or 568-conjugated anti-rabbit, anti-mouse or anti-rat IgG secondary antibody (1/500 dilution, Life technologies, CA, USA) for 2 days at 4 °C. The slides were mounted in Prolong Antifade Medium containing DAPI (Molecular Probe, MA, USA).

Ovarian cell suspensions were fixed in 1% paraformaldehyde in PBS (−) for 1 hr. Cells were washed PBS (−), and incubated overnight at room temperature with blocking solution. Immunofluorescence analysis of respectively proteins in ovarian cells conducted using the following primary antibodies: anti-γH2AX (1/200 dilution, ab11174, Abcam, Cambridge, UK), anti-SCP3 (1/500 dilution, ab97672, Abcam, Cambridge, UK) and anti-SCP1 (1/500 dilution, ab15090, Abcam, Cambridge, UK). Cells were incubated overnight at room temperature with primary antibody, and then washed and incubated with Alexa Fluor 488- or 568-conjugated anti-rabbit or anti-mouse IgG secondary antibody (1/500 dilution, Life technologies, CA, USA) at room temperature for 1 hr. The slides were mounted in Prolong Antifade Medium containing DAPI (Molecular Probe, MA, USA). Fluorescent signals were detected and quantified under the LSM710 confocal microscopy (Zeiss, Oberkohen, FRG). For all the observation, exposure time was kept the same between XX and XY female samples.

### Data from other source

Previously published RNA-Seq data from mouse E13.5 female and male PGCs (DRA003597), and P1 spermatogonial stem cells (DRA002477) were downloaded from DNA Data Bank of Japan (DDBJ), and aligned to allow comparison with the present data.

### Accession number

The RNA-Seq data from this study have been deposited in the DDBJ under the accession number DRA005345 and DRA005822.

## Acknowledgments

We thank Hidehiko Ogawa, Yayoi Obata, Hisato Kobayashi for helpful comments, and Asuka Kamio, Takumi Yoshioka for assistance of NGS data collection and data analysis.

## Author Contributions

T. Kono and A. Sakashita conceived and designed the experiments. T. Wakai provided instructions regarding the construction of the CRISPR/Cas9 targeting plasmid. A. Sakashita, C. Nishimura and Y. Sotomaru performed microinjection experiments, and carried out the phenotypic analysis of generated XY^*sry−*^-females. A. Sakashita and Y. Kawabata performed RNA-Seq and data analysis. T. Kono and A. Sakashita wrote the paper. All authors discussed the results and commented on the manuscript.

